# Anaerobic methanotrophic communities thrive in deep submarine permafrost

**DOI:** 10.1101/181891

**Authors:** Matthias Winkel, Julia Mitzscherling, Pier P. Overduin, Fabian Horn, Maria Winterfeld, Ruud Rijkers, Mikhail N. Grigoriev, Christian Knoblauch, Kai Mangelsdorf, Dirk Wagner, Susanne Liebner

## Abstract

Thawing submarine permafrost is a source of methane to the subsurface biosphere. Methane oxidation in submarine permafrost sediments has been proposed, but the responsible microorganisms remain uncharacterized. We analyzed archaeal communities and identified distinct anaerobic methanotrophic (ANME-2a/b, ANME-2d) assemblages in frozen and completely thawed submarine permafrost sediments. Besides archaea potentially involved in AOM we found a large diversity of archaea mainly belonging to *Bathyarchaeota*, *Thaumarchaeota*, and *Euryarchaeota*. Methane concentrations and δ^13^C-methane signatures distinguish horizons of potential anaerobic oxidation of methane (AOM) coupled either to sulfate reduction in a sulfate-methane transition zone (SMTZ) or to the reduction of other electron acceptors, such as iron, manganese or nitrate. Analysis of functional marker genes (*mcrA*) and fluorescence *in situ* hybridization (FISH) corroborate AOM communities in submarine permafrost sediments potentially active at low temperatures. Extrapolating potential AOM rates, when scaled to the total area of expected submarine permafrost thaw, reveals that methane could be consumed at rates between 8 and 120 Tg C per year, which is comparable to other AOM habitats such as seeps, continental SMTZ and wetlands. We thus propose that AOM is active where submarine permafrost thaws and needs to be accounted for in global methane budgets.

## Introduction

Terrestrial permafrost landscapes, which developed during glacial cold periods, are known to be a large reservoir of organic carbon (∼1300 Pg) (1). Permafrost thaw and the following microbial production of methane and carbon dioxide from liberated organic matter may act as a positive feedback to climate warming (2). The global warming potential of methane is 34 times higher than carbon dioxide over a 100 year period (3). The amount and the release rates of methane are not well constrained although they are critical for evaluating future climate change.

Several Arctic sources of methane have been identified, including methane bursts during tundra freezing (4), thermokarst lakes (5), lakes and ponds (6), wetlands (7), gas hydrates (8) and submarine permafrost (9). Submarine permafrost is much more susceptible to thawing than permafrost on land, because of overlying warm marine water causing diffusion of salt water into the sediments from the top, and because of the geothermal heat fluxes from below (9). Submarine permafrost on continental shelves of the Arctic Ocean was formed due to inundation of terrestrial permafrost by sea water during the Holocene marine transgression (10, 11). Our knowledge of carbon pools and carbon turnover in submarine permafrost is however scarce. Coastal submarine permafrost has been frozen under subaerial terrestrial conditions in alluvial/fluvial settings and has remained frozen since then (12, 13). Based on thermal modeling of permafrost development over glacial/interglacial cycles, submarine permafrost is likely to have persisted at depths for hundreds of millennia (14). Over those millennial timescales submarine permafrost reaches its freezing point, between −2° and −1°C, given saline sediment layers, after which it slowly starts degrading (10, 15).

Arctic submarine permafrost regions have the potential to emit large amounts of methane to the atmosphere (16), however the fluxes are not well constrained. Estimates of methane release of 8.0 Tg y^-1^ from the East Siberian Arctic Shelf (comprising Laptev Sea, East Siberian Sea and the Russian part of the Chuckchi Sea) equal the release of methane from the entire world ocean (9). In contrast, recent estimates for the Laptev and East Siberian Seas are almost three times lower (2.9 Tg y^-1^) (17). Methane is mainly released from the sediment but the source is unclear (18). It is expected that permafrost degradation creates pathways for the release of gas captured within submarine permafrost sediment or within gas hydrates underneath (16). Low δ^13^C-CH_4_ values indicate a biogenic origin of methane released from submarine permafrost (13, 19). A drop of methane concentrations at the permafrost thaw front deep inside the sediments, which coincided with increasing δ^13^C-CH_4_ values from −70‰ to −35‰ indicated anaerobic oxidation of methane (AOM) (13). Sulfate penetrating from the seabed into the sediment created a deep sulfate-methane transition zone (SMTZ) similar to other geological formations in the subsurface (20) indicating that sulfate is potentially the electron acceptor for AOM (21).

Most studies on AOM in the marine system have focused on ANME archaea that couple the oxidation of methane to the reduction of sulfate via a syntrophic lifestyle with sulfate-reducing bacteria (SRB) (22–24). Three main marine clades, ANME-1a/b, ANME-2a/b/c, and ANME-3 have been identified so far (25). These clades are associated with sulfate-reducing bacteria of the genera *Desulfosarcina*, *Desulfococcus*, *Desulfobulbus* and *Desulfofervidus*, which belong to the class *Deltaproteobacteria* (25) and of the phylum *Thermodesulfobacteria* (26). Besides the marine ANMEs there is evidence of terrestrial AOM driven by the ANME-2d clade (27, 28). ANME-2d sequences were detected in wetland and permafrost habitats (29–31) so they might mitigate the release of methane in these enviroments. A recent study on wetlands showed that even low amounts of sulfate (μM) are sufficient to couple AOM with sulfate reduction in terrestrial environments, although the identification of involved microorganisms is missing (32). Alternative electron acceptors for AOM such as iron (33, 34), manganese (33) as well as nitrate (27, 35) and humic substances (36, 37) have been suggested in some marine and freshwater studies. To date, the microbial mitigation of methane emissions from permafrost environments is only evident in aerobic soils and sediments (38, 39), while communities potentially anaerobically oxidizing methane in thawing permafrost in the deep subsurface remain unexplored (8). We hypothesize that active marine and terrestrial ANME clades exist in thawing submarine permafrost similar to microorganisms found in other shallower marine SMTZ (40, 41) or in terrestrial enrichments (27, 35), and that these clades function as an efficient methane filter for a diffusive methane release.

We investigated AOM in two deep submarine permafrost cores (48 m and 71 m) with different stages of submarine permafrost thaw (42) that were inundated for about 540 and 2500 years, respectively (11, 13). To identify microbial assemblages involved in AOM, we performed high-throughput sequencing of archaeal and bacterial 16S rRNA genes and of the functional marker gene methyl co-enzyme reductase (*mcrA*) that is encoded by both methanogens and anaerobic methanotrophs. We quantified *mcrA* and dissimilatory sulfate reductase (*dsrB*) gene marker by quantitative PCR (qPCR) and correlated their abundance with the 16S rRNA gene abundance of AOM-related microorganisms. We visualized AOM-related microorganisms by catalyzed reporter deposition fluorescence *in situ* hybridization (CARD-FISH) and constructed ANME-specific 16S rRNA gene clone libraries to resolve their diversity in submarine permafrost sediments. Pore water geochemistry i. e., concentrations of methane, nitrate, sulfate, iron, and manganese was analyzed to verify potential horizons of methane consumption. Analysis of stable carbon isotope geochemistry of methane and microbial membrane lipid biomarker were performed to evaluate the activity of AOM communities *in situ*.

## Results

### Isotopic evidence of AOM and potential electron acceptors

Methane concentrations were low in the ice-bonded permafrost section of the Cape Mamontov Klyk core (hereafter C2) inundated for ∼2500 years (13) and peaked at approximately 52 m below seafloor (bsf) with a concentration of 284 μM (Fig. 1A). In contrast, methane concentrations in the ice-bonded permafrost section of the Buor Khaya core (hereafter BK2) were on average 38 times higher than in C2 (Fig. 2A). δ^13^C values of methane in the ice-bonded permafrost showed values between −71 and −53‰ typical for a biological origin (*43*). Above the ice-bonded permafrost, δ^13^C values of methane ranged between −37 and −30‰ within a few meters on top of the thaw front and stayed constant to a depth of about 12 mbsf (13). Above 12 mbsf δ^13^C-values of methane showed high dynamics with values ranging between −67 to −32‰. Overall methane concentration and δ^13^C values in both cores revealed opposing dynamics indicating both, microbial production and consumption of methane.

**Fig. 1.**
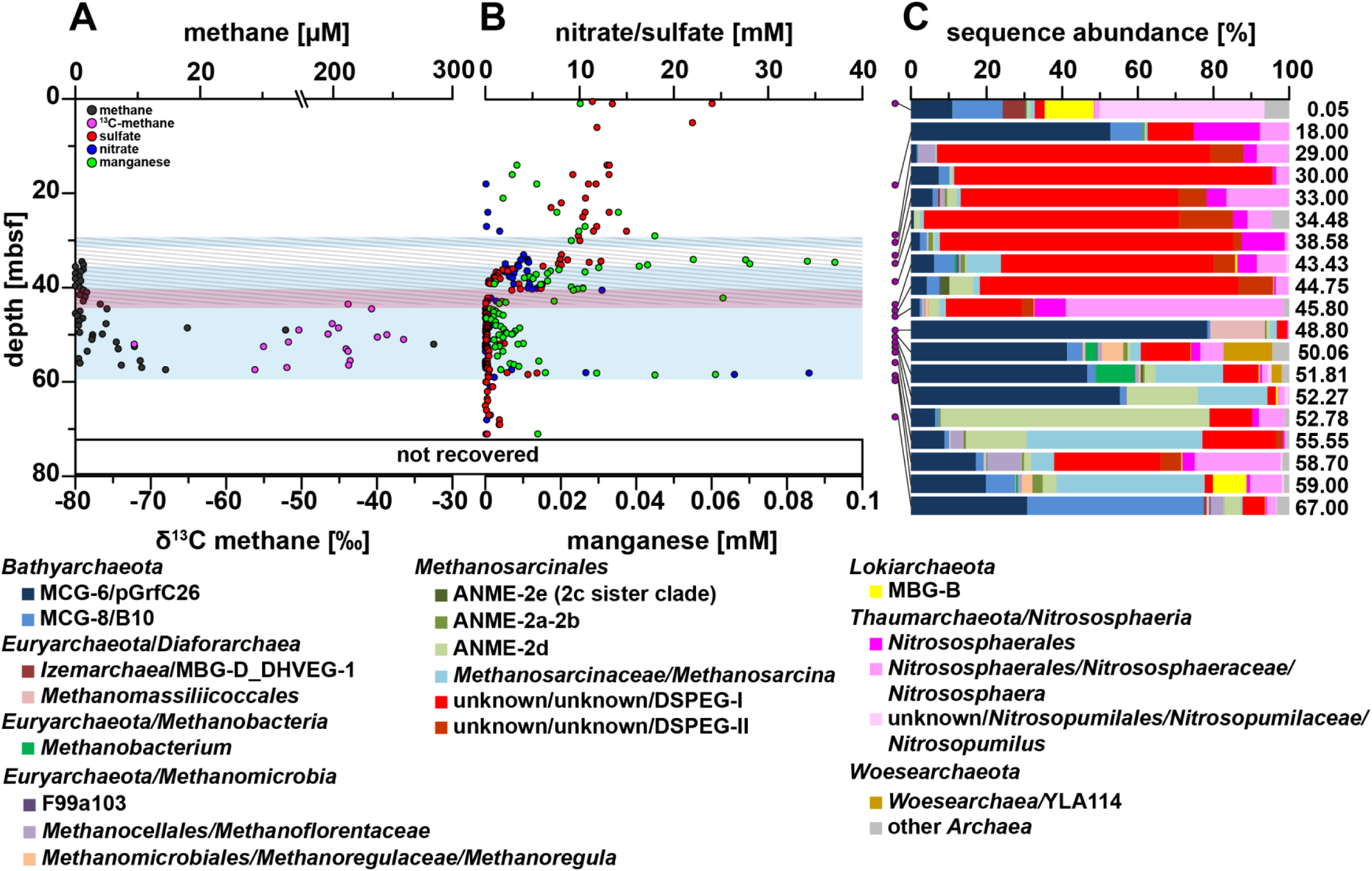
Pore water profiles of methane, nitrate, sulfate, manganese and relative archaeal abundance in the Mamontov Klyk core C2. (A) Methane concentrations are shown in black and corresponding δ^13^C-values in magenta (*19*). (B) Nitrate concentrations are shown in blue, sulfate concentrations in red, and manganese concentrations in green. (C) Purple dots on the left shows depth location in the core. Relative abundance of archaeal sequences are shown as bar plots. Colors of bars refer to the taxa in the legend below figure. Numbers on the left refer to the exact depth in the core. ANME= ANaerobic MEthanotrophic archaea, DHVEG-1= Deep Hydrothermal Vent Euryarchaeotal Group 1, DSPEG= Deep Submarine Permafrost Euryarchaeotal Group, MBG-B= Marine Benthic Group B, MBG-D= Marine Benthic Group D, MCG= Miscellaneous Crenarchaeotal Group. The shaded area represents the permafrost degradation zone, the red area a sulfate-methane transition zone, and the light blue areas ice-bonded permafrost.

**Fig. 2.**
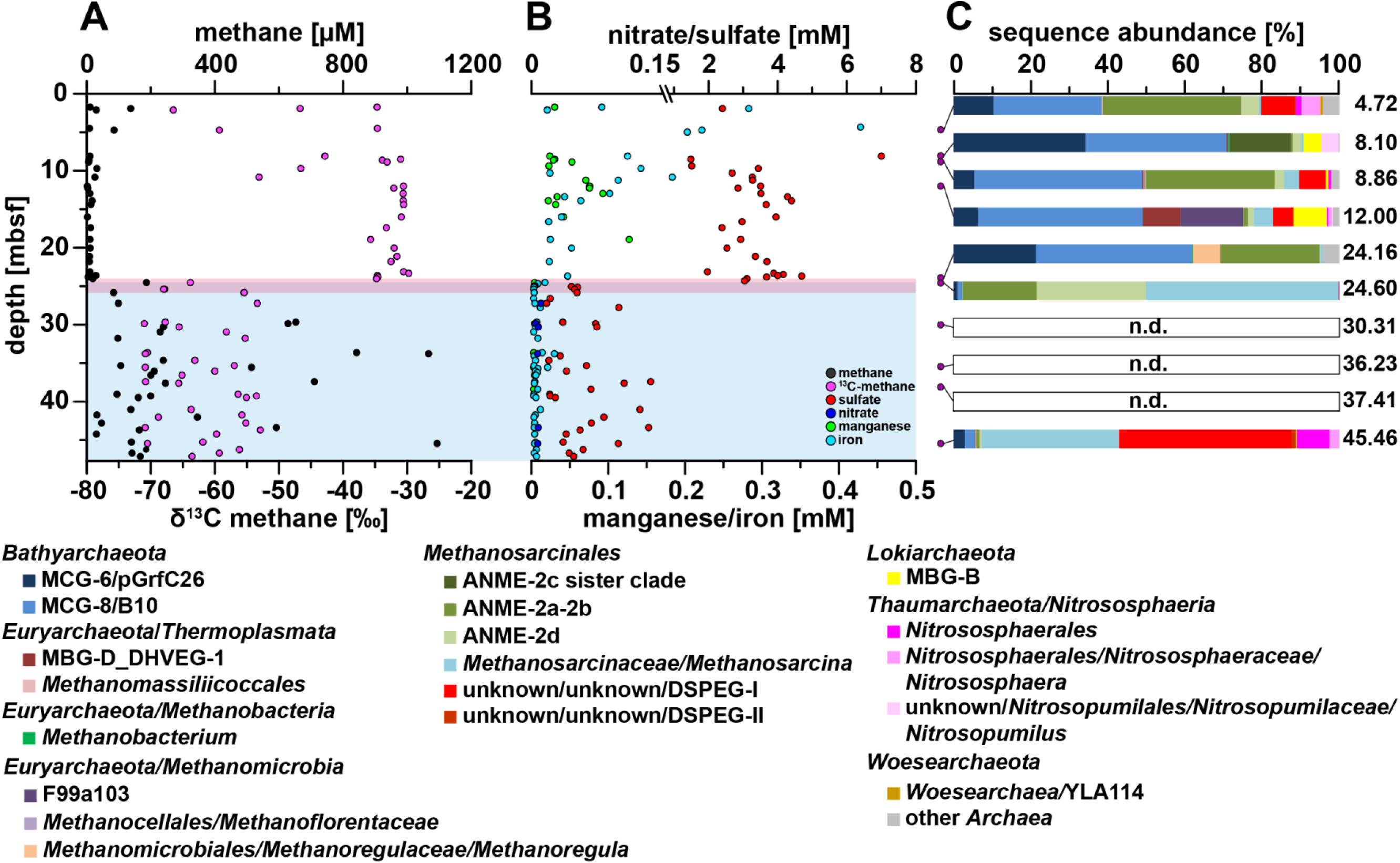
Pore water profiles of methane, nitrate, sulfate, iron, manganese and relative archaeal abundance in the Buor Khaya core BK2. (A) Methane concentrations are shown in black and corresponding δ13C-values in magenta (19). (B) Nitrate concentrations are shown in blue, sulfate concentrations in red, manganese concentrations in green, and iron concentrations in light blue. (C) Purple dots on the left shows depth location in the core. Relative abundance of archaeal sequences are shown as bar plots. Colors of bars refer to the taxa in the legend below figure. Numbers on the left refer to the exact depth in the core. ANME= ANaerobic MEthanotrophic archaea, DHVEG-1= Deep Hydrothermal Vent Euryarchaeotal Group 1, DSPEG= Deep Submarine Permafrost Euryarchaeotal Group, MBG-B= Marine Benthic Group B, MBG-D= Marine Benthic Group D, MCG= Miscellaneous Crenarchaeotal Group. The red area represents a sulfate-methane transition zone and the light blue area ice-bonded permafrost.

Estimated release rates of methane from the ice-bonded permafrost of BK2 would yield potential AOM rates of 2.1 nmol cm^-3^ d^-1^ ± 1.2 nmol cm^-3^ d^-1^ (13).

Sulfate penetrated into the ice-bonded permafrost section of core C2 and into the thaw front of core BK2 indicating the presence of a SMTZ (Fig. 1A, B and 2A, B). Sulfate concentrations of 12 to 24 mM in the upper layers of core C2 were caused by the penetration of marine water into the sea bed. Sulfate concentrations decreased with depth and penetrated almost 10 m into the ice-bonded permafrost. Here, sulfate concentrations of up to 12.5 mM were still comparably high with up to 12.5 mM. Sulfate in the marine affected layers of BK2 showed lower concentrations of 2 to 7 mM (Fig. 2B). In core C2, other potential terminal electron acceptors (TEA) such as nitrate, manganese, and iron were detected in the permafrost sediment layers (Fig. 1B, C). Nitrate and manganese concentrations displayed high fluctuations in ice-bonded layers showing marine influence (42) in the sediment i.e. between 26 and 42 mbsf. A high peak of nitrate and manganese was also observed at the lower boundary of the ice-bonded permafrost (58.5 mbsf). In the ice-bonded permafrost part of BK2, manganese, iron and nitrate were close to or even below the detection limit. In the unfrozen part, manganese and iron concentrations were highly variable with concentration an order of magnitude higher (Fig. 2B, C).

### Archaeal community structures of degrading permafrost

The upper layer core C2, inundated since 2,500 years, was dominated by sequences of typical marine archaeal taxa. These were represented by aerobic ammonia-oxidizing *Thaumarchaeota* of the genus *Nitrosopumilus* (∼44%), *Lokiarchaeota* found at hydrothermal vents (∼15%), by the marine benthic group D (6%) and other marine taxa such as the marine group II (∼2%, Fig. 1C). At 18 mbsf *Bathyarchaeota* sequences of the MCG-6/pGrfC26 clade dominated (∼53%) the archaeal community. This clade is known to degrade polymeric carbohydrates in the sulfate reduction zone (44). In same depth a shift towards taxa that are commonly found in terrestrial environments such as the ammonia-oxidizing *Nitrososphaerales* (∼25%) was observed. Here we also detected a so far unknown clade, which we designated as **D**eep **S**ubmarine **P**ermafrost **E**uryarchaeotal **G**roup I (DSPEG I, ∼12%). This new clade is closely related to *Methanomassiliicoccales* of the superclass *Diaforarchaea.* DSPEG I increased in relative sequence abundance with depth (Fig. 1C and 3). DSPEG I and a sister clade (designated DSPEG II) were most dominant in the sediment horizons between 29 and 44.8 mbsf, where they represented between 62 to 84% of all archaeal sequences (Fig. 1C). Both groups showed less than 82% similarity values to the next cultured representative *Methanomassiliicoccus luminyensis* (*45*) that places them on a new order level. Other abundant sequences belong to *Methanoflorentaceae* (∼0.1-5%), ANME-2d (∼0.2-6%), *Methanosarcina* (∼0.2-9%), *Bathyarchaeota* MCG-6 and MCG-8 (∼0.8-11%), and *Nitrososphaerales* (∼3.7-22%). At 45 mbsf there was a pronounced shift towards *Nitrososphaerales* sequences (∼65%). In the sediment layers between 49-52 mbsf, where most of the electron acceptors showed low concentrations, *Bathyarchaeota* of the MCG-6/pGrfC26 clade dominated (∼41-78%). In these sediment horizons, many methanogen-related sequences appeared, such as *Methanoregula* (∼0.1-6% between 50.06-52.3 mbsf), *Methanosarcina* (∼2-18%), *Methanobacterium* (∼3-10% 50.1-52.3 mbsf) and *Methanomassiliicoccaceae* (0.7-14% between 48.8-50.6 mbsf). Between 50.1 and 51.8 mbsf *Woesearchaeota* of the YLA114 clade were present (∼3-13%). In core C2 ANME-2d sequences increased in relative abundance from less than 1% at 48.80 mbsf with less than 1% to 18% at 52.3 mbsf and up to 70% at 52.8 mbsf. The high abundance of anaerobic methane-oxidizers coincides around the highest peak of the methane concentration profile in the ice-bonded permafrost (284 μM) at 52 mbsf (Fig. 1). In the same depth, about 1.1% of sequences are related to *Methanosarcinaceae*/ANME-3 (*24*). A second, much smaller peak of the methane concentration profile (10 μM) at 55.6 mbsf still showed 16% ANME-2d sequences. At the same depth *Methanosarcina* represented the most abundant group (∼47% of all archaeal sequences) and DSPEG I and II related sequences were the second most abundant (∼21%, Fig. 1C). Towards the lower boundary of the ice-bonded permafrost at 58.7 mbsf, the DSPEG I and II increased again in sequence abundance (∼33%). The sediment horizon directly underneath the permafrost contained some sequences related to marine environments, such as *Lokiarchaeota* (∼9%), and ANME-2a/b (2.5%). The deepest sediment sample showed mostly *Bathyarchaeota* sequences of subgroups MCG-6 and MCG-8 (∼77%) and DSPEG I and II, ANME-2d, *Methanoflorentaceae* and *Nitrososphaerales* (∼6%, 4%, 3% and 2%, respectively) sequences.

The more recently inundated core BK2 contained a mixture of marine and terrestrial archaeal sequences in the depth between 4.7 and 12 mbsf. Marine clades were represented by ANME-2a/2b (∼0.5-35%), *Methanomicrobia* of clade F99a103 (∼0.1-16%) that were first discovered at a hydrothermal chimney, ANME-2c sister clade (∼16%, 8.1 mbsf), Marine Benthic Group D (∼0.1- 10%), *Lokiarchaeota* (∼0.7-9), and *Nitrosopumilales* (∼0.3-4%). Terrestrial clades were represented by *Methanosarcinales/Methanosarcina* (∼0.7-15%), the DSPEG I and II (∼0.1-9%), and *Nitrososphaerales* (∼0.2-5%). The high numbers of ANME-2a/b sequences (Fig. 2D) at depths of ∼5 and ∼9 mbsf correlated with highest δ^13^C-methane values in the uppermost 12 m that were characterized by highly variable methane δ^13^C-vcalues (−67 to −31‰, Fig. 2D). Nevertheless, *Bathyarchaeota*-related sequences of the clades MCG-6 and MCG-8 (∼40-71%) dominated throughout the layers above the ice-bonded permafrost. In contrast, they drastically decreased (∼2%) in the upper layer of the ice-bonded permafrost. The sediment horizon between 24.0 and 24.7 mbsf was characterized by opposing gradients of sulfate and methane. In the upper part of this layer that is strongly marine influenced ANME-2a/b made up about 26% of all archaeal sequences while in the lower and still partially ice-bonded part of the permafrost thaw front both ANME-2a/b and ANME-2d clades were detected in high abundances (about 19% and 28% respectively (Fig. 2C). In the ice-bonded permafrost of core BK2, archaea were only detected in about 45.46 mbsf. Here, *Methanosarcina* (∼37%) and DSPEG I and II (∼46%) related sequences dominated. A detailed description of the bacterial community structure and diversity of both cores was described elsewhere (42).

### Diversity of ANME-related groups in degrading permafrost

Both cores displayed layers where ANME sequences dominated the archaeal community. The major two groups, ANME-2a/b and ANME-2d showed a remarkably high diversity with 23 and 17 operational taxonomic units (OTUs), respectively (Fig. 3 and 4A). ANME-2d separated into 4 different clusters (Fig. 4B), while the majority of all sequences fell into cluster 1. This cluster also contained *Candidatus* Methanoperedens nitroreducens BLZ1 (46), known to either use nitrate (27) or iron (47) as electron acceptor for methane oxidation. Besides the two dominant ANME-groups, we also detected sequences in a sister clade of ANME-2c (designated ANME-2e) with 5 OTUs (Fig. 3 and 4 for details see also Fig. S1). Other sequences of this group were retrieved from cold seep sites and mud volcanoes. Moreover, we detected 1 OTU that fell into the ANME-3 cluster (Fig. 3 and 4, for details see also Fig. S2). Clone libraries of the SMTZ, with specific ANME-2a primers revealed mainly ANME-2a/b sequences and a few ANME-2d as well as one sequence of the ANME-2c sister clade (Fig. 4A, for details see also Fig. S1). Corroborative FISH analysis of the SMTZ detected ANME-2a consortia with the specific ANME-2a-647 probe, while detection with a specific probe for *Desulfosarcina/Desulfococcus* showed no signal in any consortium (Fig. S3).

**Fig. 3.**
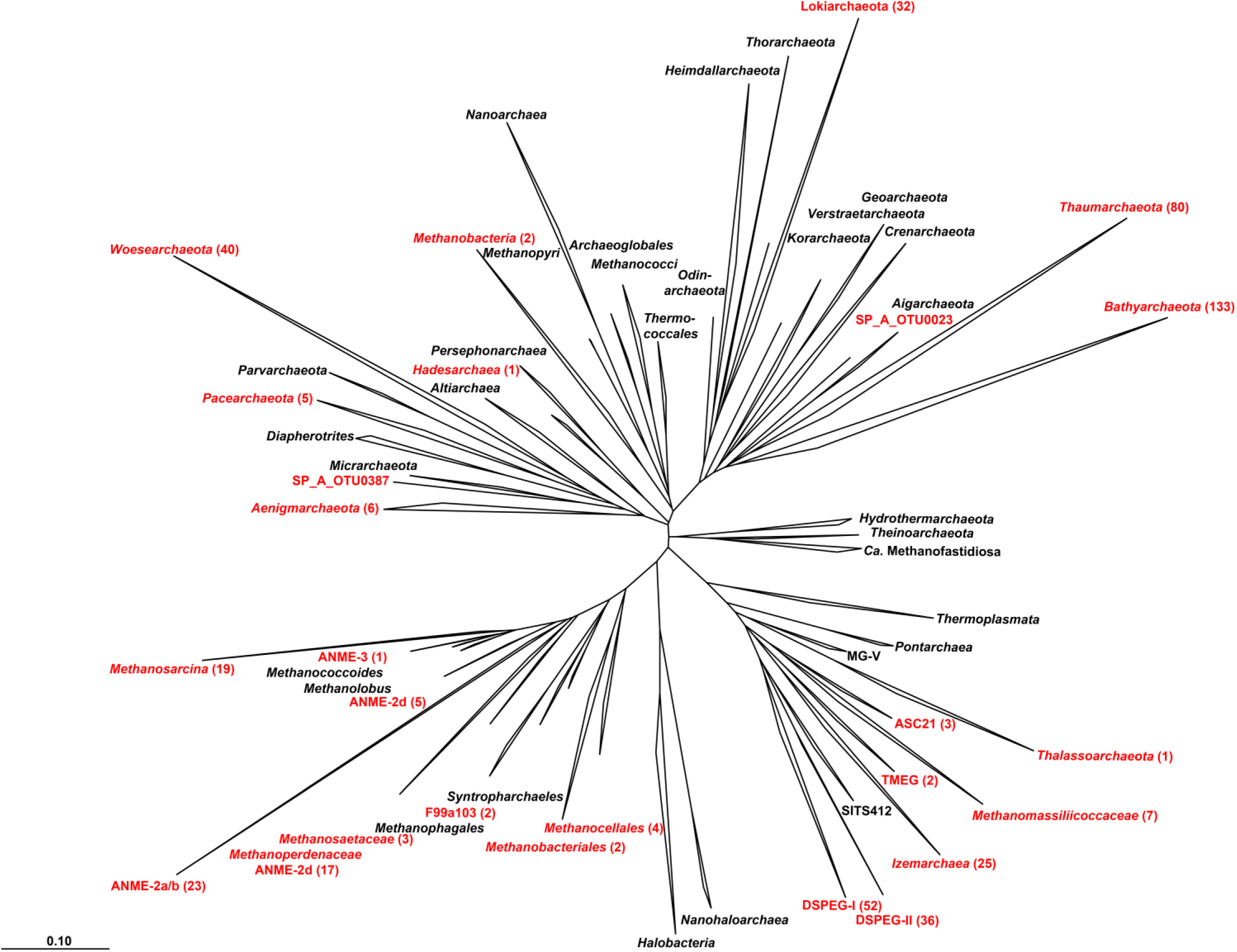
Phylogenetic affiliation of submarine Permafrost archaeal sequences based on 16S rRNA gene. Taxonomic cluster in red contain sequences of submarine permafrost. Numbers in brackets show the number of OTUs per cluster. The scale bar represents 10 percent sequence divergence. The tree was rooted with the DPANN superphlyum.

**Fig. 4.**
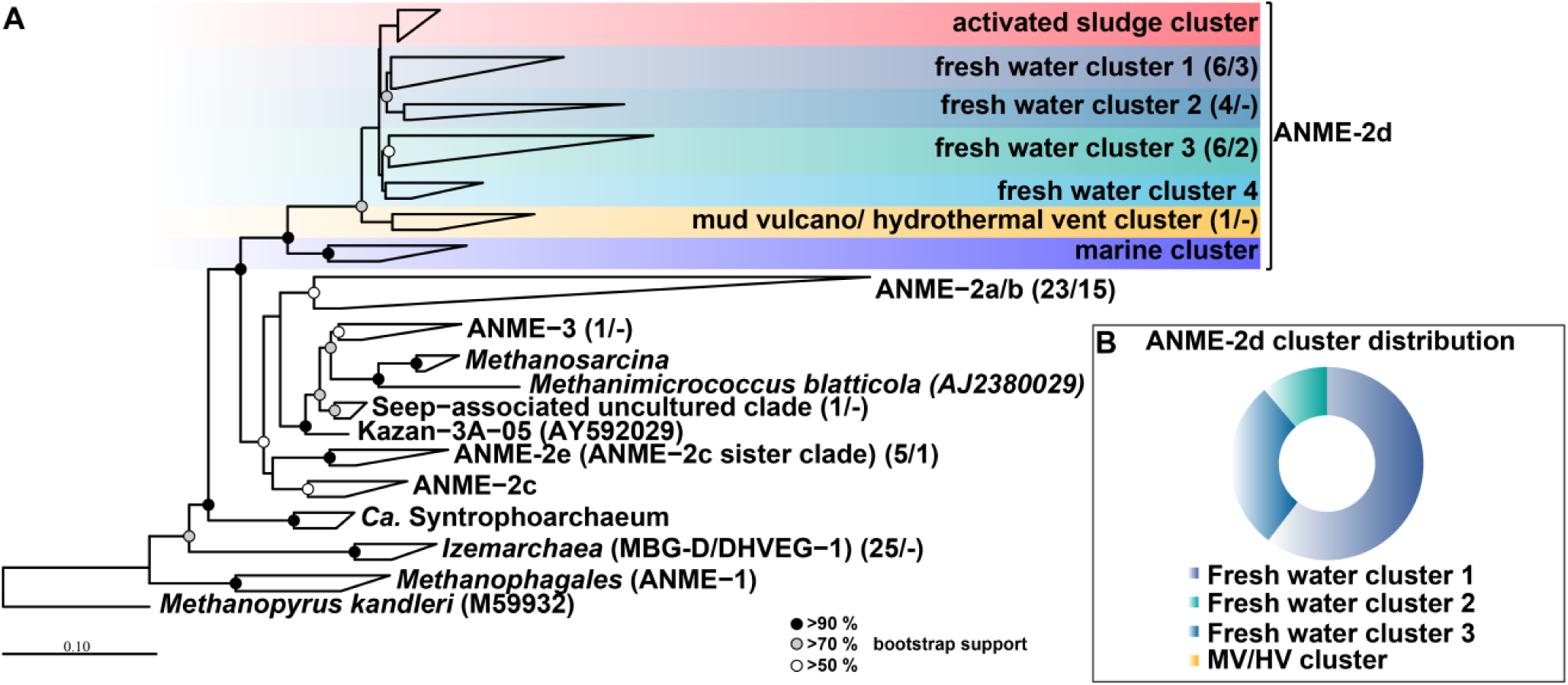
Phylogenetic affiliation of ANME sequences based on 16S rRNA gene. (4A) The first number in the parenthesis represents the number of OTUs from the Illumina MiSeq sequencing, while the second number represents the number of sequences of the clone library with ANME-specific primers. Methanopyrus kandleri was used as outgroup. The scale bar represents 10 percent sequence divergence. The inlet round ring diagram represents the whole relative distribution of all ANME-2d sequences in both submarine permafrost cores (4B).

Canonical correspondence analyses (CCA) illustrated that ANME-2a/b and ANME-2e cluster with other marine-related archaea and are mainly influenced by salinity, iron and sulfate (Fig. 5). Spearman rank correlations revealed that the abundance of ANME-2a/b significantly correlated with iron concentrations (R=0.65, *p*< 0.03). ANME-2d sequences cluster together with sequences of *Methanosarcina* and *Bathyarchaeota* of the MCG-6/pGrfC26 clade, while only the latter correlate with methane concentrations (Spearman correlation R=0.41, *p*> 0.09,).

**Fig. 5.**
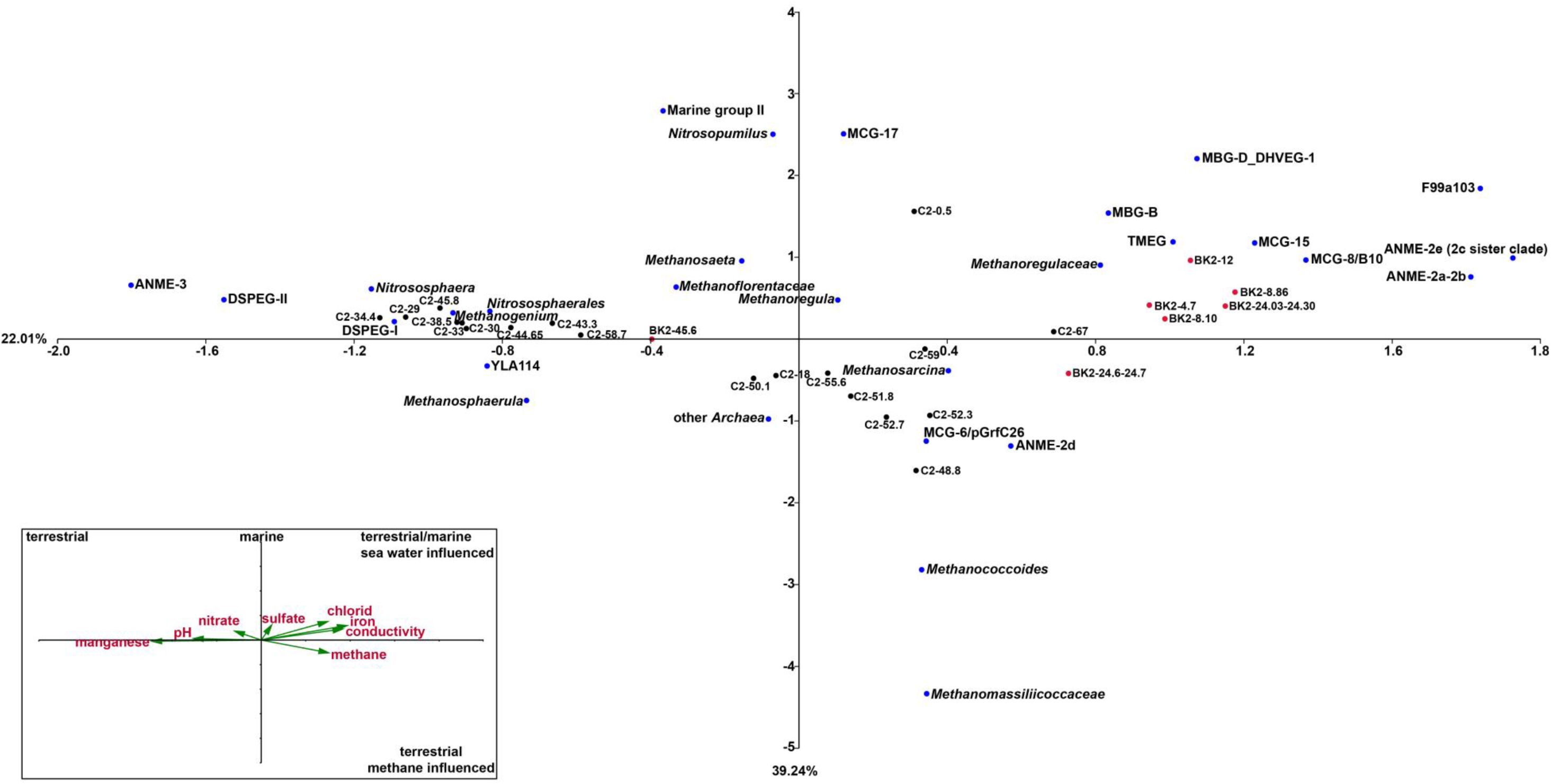
Canonical correspondence analysis (CCA) of environmental factors and archaeal taxa that contributed more than 1% in any of the depth. Inlet: Environmental factors are plotted as triplot with scaling 1.5 shown as black arrows and red terms. Samples of core C2 are projected as black dots and samples of BK2 as red dots. The archaeal taxa are shown as blue dots. Percentages at axes represent the ‘Eigenvalues’ that explain the variability for the first two axe

### Functional groups responsible for methane cycling

Quantitative detection of the methyl co-enzyme reductase subunit A (*mcrA*) as functional marker for methane production (methanogens) and AOM (ANME) showed a decrease with depth in the ice-free permafrost of both cores (0 to 29 mbsf for C2, and 0 to 24 mbsf for BK2, respectively, Fig. S4A and S4B). In the ice-bonded permafrost of core C2 we detected an increase in *mcrA* copy numbers with the highest copy numbers g^-1^ wet weight (0.3 – 1.2 x 10^5^ ± 0.4 - × 10^3^) between ∼45 and 52 mbsf. At 53 mbsf total *mcrA* copy numbers decreased, while 16S rRNA gene sequences at the same depth were almost exclusively affiliated with ANME-2d (Fig. 1C and Fig. S4A). 454 amplicon analyses of the *mcrA* sequences showed a relative increase of ANME-2d-related *mcrA* genes towards the layers with highest ANME-2d-related 16S rRNA gene sequences between 52.3 and 55.6 mbsf (Fig. 2C and Fig. S5). Quantification of ANME-2d with recently designed *mcrA* primers in the ice-bonded permafrost (48) showed up to 1.2 ± 0.5 x 10^5^ copy numbers g^-1^ wet weight at approximately 57 mbsf and an average copy number of 5.1 ± 1.5 x 10^4^ g^-1^ wet weight over all depth (Fig. S6).

Analysis of the dissimilatory sulfate reductase subunit B (*dsrB*) for SRB mainly followed the depth distribution of *Desulfosporosinus* 16S rRNA sequences in the C2 core, with a few exceptions (29.0, 30.0, 34.4 and 38.5 mbsf) where an increase in the *dsrB* could not be related to any known sulfate-reducers (Fig. S7A).

*McrA* copy numbers in the BK2 slightly increased at the SMTZ and may be underestimated due to a primer bias against ANME *mcrA* genes (48), while 16S rRNA gene sequences showed 25-47 % ANME-related sequences at the SMTZ (Fig. S4B). The *dsrB* copy numbers slightly increased at the SMTZ, although SRB represented < 1% of the total bacterial 16S rRNA gene sequences (Fig. S7B). Phylogenetic reconstruction of SRB at the SMTZ indicated that all sequences of the upper part belonged to typical marine clades SEEP-SRB1 and another seep-associated *Desulfobacterium anilini*-group (red OTUs, Fig. S8). In the lower part of the SMTZ that is dominated by ANME-2d sequences, seep-associated SRB were below detection limit, while only *Desulfosporosinus*-related sequences (blue OTU, Fig. S8) could be observed. Still, SRB at the thaw front represented < 1% of bacterial sequences.

An attempt to analyze lipid biomarker revealed very low concentrations that prevented a precise isotopic analysis to conclude activity of AOM microbial communities. Nevertheless, calculated branched and isoprenoid tetraether (BIT) indices showed values of terrestrial origin between 0.99 to 1 (49). Moreover calculated methane indices (MI) for isoprenoid GDGTs showed values between 0.85 to 0.99 in the ice-free sediment layer and 1 at the SMTZ (50). The ratio of archaeal to bacterial ether lipids showed dominant archaeal values in the SMTZ (Fig. S9). For further discussion on lipid biomarker see supplementary material.

## Discussion

Since the last decade methane concentrations in the atmosphere have been increasing again, but the sources and mechanisms are not fully understood (51). One potentially large but highly controversial source of methane is submarine permafrost, which faces drastic changes due to global temperature increase and associated Arctic sea ice reduction (52). Our study provides multiple lines of evidence that ANME communities are present in submarine permafrost layers where methane got consumed. We thus propose the microbial mitigation of methane release from thawing deep submarine permafrost on the Siberian Arctic Shelf. We detected both marine and terrestrial ANME clades likely involved in the AOM process at various depths, not only at the permafrost thaw front, but also in still ice-bonded permafrost that undergoes degradation. Our study indicates that AOM occurs at temperatures around 0°C.

Pore water methane concentrations in the two cores were in the typical range of deep sediments and soils (20, 53). The permafrost thaw front at the BK2 site, inundated about 540 years ago revealed a deep SMTZ. Here, the large shift of the methane δ^13^C-values can only be explained by microbial oxidation. The calculated potential rates (2.1 nmol cm^-3^ d^-1^± 1.2 nmol cm^-3^ d^-^1) (*13*) are typical for marginal SMTZ and exceed those of subsurface SMTZ (*25*). At the permafrost thaw front of BK2, we detected marine and terrestrial ANME clades that potentially mitigate the methane release into overlying sediment layers. The upper ice-free part of the SMTZ represents a typical marine sulfate-dependent AOM community of ANME-2a/b (Fig. 2C) affiliated with SEEP-SRB1/seep-associated *Desulfobacterium anilini*-group (Fig. S7A). These sulfate-dependent AOM communities are of marine origin and are often found in mud vulcanoes, methane and hydrocarbon seeps (54). We could visualize ANME-2a *in situ* using CARD-FISH further supporting their occurrence at the permafrost thaw front of BK2. The molecular detection of associated SRB are, however, not conclusive. This might be due to SEEP-SRB partners that are not targeted by the DSS685 probe (55) or by yet unidentified bacterial partners (56). The lower part of the SMTZ was characterized by a clear transition to a terrestrial AOM community closely related to *Candidatus* Methanoperedens nitroreducens (ANME-2d) that might be capable of using alternative TEA such as nitrate and iron (27, 47). In a recent genomic study it has been proposed that these organisms have the genetic repertoire for an independent AOM process without a bacterial partner (46). The ether lipid MI values of BK2 were highest in the SMSTZ and further support a potential involvement of AOM communities (50).

The high abundances of marine-derived ANME-2a/b sequences in several depths above the SMTZ of BK2 with otherwise BIT indices towards an exclusively terrestrial origin show that terrestrial and marine sediments were mixed in the upper meters of BK2 (Fig. 2). The terrestrial sediments were thus influenced by sea water penetrating down to the permafrost table and thereby transport of marine organisms into deeper layers occurred (Fig. 2). This is consistent with profiles of other environmental parameters (pH, temperature, isotopes and ion concentrations) and the stratification and composition of the bacterial communities (42).

In core C2 the layer between 10 and 29 mbsf is characterized by alternating frozen and partly thawed sediments (Fig. 1) and showed high fluctuations in the concentrations of nitrate and manganese. These fluctuations indicate degradation of thawing permafrost (shaded area, Fig. 1) and an increase in microbial activity resulting in the consumption of labile carbon pools (*57*). Active processes in ice-bonded permafrost close to thawing (mean temperature: −1.2 ± 0.2 °C, table S1) can take place in liquid water films surrounding mineral particles, which form a network in which microbial activity is expected (58). Microbial activity may thereby be supported through sulfate as additional electron acceptor (59) penetrating into the ice-bonded sediment. Indeed, sulfate concentrations decreased and *dsrB* copy numbers increased with depth pointing towards active sulfate reduction and effective anaerobic organic matter decomposition (60). In C2, the SMTZ occurred below the actual permafrost thaw front but inside the ice-bonded permafrost and was characterized by high manganese and nitrate concentration. This could promote AOM with alternative TEA (33, 35) and shows ongoing degradation of ice-bonded permafrost. Also, the detection of low abundances of ANME-2a/b in this layer shows their migration into permafrost by downward marine water intrusion (42). Unlike in core BK2, the ice-bonded layer of C2 was almost absent of methane. At the same time ANME-2d occurred almost entirely throughout this layer. The relatively low copy numbers of ANME-2d detected with specific mcrA primers and compared with total cell counts (42) are still in the range of North Sea and River sediment (48). The highest abundance of ANME-2d coincided around the highest methane concentration, which we consistently detected by several molecular approaches (16S rRNA, general *mcrA* and ANME-2d-specific *mcrA*). Taken all this together we suggest that ANME-2d organisms are responsible for AOM in the ice-bonded permafrost before it thaws. An alternative explanation for the low methane concentrations and the low abundance of ANMEs in most of the ice-bonded layers are a relic of AOM communities that were active under different environmental conditions in the past. Whether the methane was trapped in the permafrost during freezing or it was produced by microbial degradation of organic matter under recent *in situ* conditions cannot be resolved since radiocarbon analysis would give similar results in both cases.

Even though sulfate reduction might be relevant for organic matter mineralization (60), links to sulfate-dependent AOM were not observed in the core C2. While C2 exhibited high copy numbers of *dsrB* and of SRB-related sequences, sulfate reducers were almost exclusively linked to *Desulfosporosinus* that have not been observed in AOM consortia so far. This genus belongs to the phylum *Firmicutes* and has been found in natural terrestrial environments such as peatlands, aquifer and permafrost (61, 62). Other TEA than sulfate, such as nitrate, iron and manganese, could also be related to AOM, and showed relatively low concentrations at the highest occurrence of ANME-2d sequences at 52 mbsf. This serves further as an indication of methane consumption during the process of AOM as known from physiological studies of ANME-2d enrichments (27, 47) but direct evidence is missing. Finally, besides the detected electron acceptors, other TEA such as humic acids could serve in the AOM process. Humic acids were shown to be involved in the AOM (36, 37) in peatlands where they were detected in high concentrations. Humic acids are produced during organic matter degradation and soil formation (63) and could thus play a role in thawing permafrost, too.

Two clades, which we named DSPEG I and DSPEG II, mainly occurred in submarine permafrost layers that showed relatively high concentrations of iron and manganese in the pore water (Fig. 1B, C). This could point towards an involvement in iron (III) and manganese (IV) reduction within the anaerobic oxidation of organic matter, in addition to sulfate reduction (*62*). Spearman rank analysis showed significant correlations between DSPEG I and DSPEG II and manganese concentration (R= 0.58, p< 0.006 and R= 0.53, p< 0.02, respectively) and negative correlations with methane (R= −0.57, p< 0.02 and R= −0.64, p< 0.004, respectively) as illustrated in the CCA (Fig. 5). We propose that these two groups reflect indicator taxa (table S2) for degrading permafrost. This is also supported by high occurrence of DSPEG I and DSPEG II (∼19 to 28%, Fig. 1C) at the lower boundary of the ice-bonded permafrost, representing bottom-up permafrost thaw. Here, high concentrations of sulfate, nitrate, and manganese show an upwardly directed thaw process (Fig. 1B). Environmental sequences affiliated with the DSPEG groups were mainly retrieved from cold environments (64, 65) and pristine aquifers (66), which further support an active role in low temperature habitats.

Taken together our molecular and biogeochemical data from two submarine permafrost cores indicate several microbial assemblages that are able to prevent the release of trapped or recently produced methane into the overlying unfrozen sediment following submarine permafrost thaw. Therefore, we challenge the assumption that high methane emissions reported for the Siberian Arctic Shelves originate from degrading submarine permafrost itself (9) and suggest different mechanisms to be responsible, such as diffusion or ebullition through discontinuities in permafrost or the release from gas hydrates (8, 67) at a limited spatial scale. Microbial assemblages in deep permafrost environments are usually associated with slow growth rates (68) and low abundance (69), and their activity is difficult to measure. New approaches such as BONCAT-FISH (56) have the potential for more direct detection of active microorganism and the analysis of their genomic potential.

Based on coarse estimates of the submarine permafrost area (∼3 million km^2^) (*2*), the potential of AOM would result in the consumption of 8 to 30 Tg C per year (lowest and highest concentration) assuming a 1 m active AOM layer or 30 to 120 Tg C per year (lowest and highest concentration) assuming a 4 m active AOM layer, as indicated in Figure 1. These estimates fit well with AOM in other environments such as wetlands 200 Tg C y^-1^ (*32*), marine SMTZ (< 50 Tg C y^-1^) and seep sites (< 10 Tg C y^-1^) (70). While in BK2 AOM at a deep SMTZ is evident, the longer inundated core C2 lacks a clear SMTZ but rather shows a patchy occurrence of possible AOM activity. Therefore, further deep drilling into submarine permafrost is necessary for a better representation of the global relevance of AOM in degrading submarine permafrost.

Our study provides first molecular evidence of microbial communities in thawing submarine permafrost that are likely involved in AOM processes. In addition, many archaeal taxa such as the newly designated DSPEG groups, a large diversity of *Bathyarchaeota*, and *Thaumarchaeota* closely related to nitrogen cycling organisms are detected. Their function is unknown and need further investment to understand their contribution in organic matter degradation of permafrost thaw processes.

## Materials and Methods

### Site description and sampling

Sediment cores were drilled along two transects from terrestrial permafrost to offshore, submarine permafrost in the Siberian Laptev Sea: Mamontov Klyk (2005) and Buor Khaya (2012) (table S1). The outermost submarine permafrost cores influenced longest by inundation were chosen for chemical and microbial investigation. The Mamontov Klyk study site was located in the western Laptev Sea (Fig. S10). The drilled core at 11.5 km offshore is characterized by three different lithostratigraphical units and contains two ice-bonded permafrost layers between 29.5 - 30.4 mbsf and 34.3 – 58.7 mbsf, respectively (12). The core had a total length of 71 m and contained sandy loam sediment with on average 0.38 % organic carbon and an average C/N ratio of 14.

The study site of Buor Khaya is located in the central Laptev Sea on the western part of the Buor Khaya peninsula (Fig. S10). The submarine core contained again 3 lithological units with an ice-bonded permafrost layer between 24.7 – 47.6 mbsf (13). The core had a total length of 47.7 m. The retrieved sediment material consisted of fine sand and contained about 1% organic carbon with a C/N ratio of 14.

The drilling was performed with a mobile drilling rig (URB-2A-2/URB-4T) and is described elsewhere (12, 13). The frozen cores were split along the vertical axis under aseptic conditions. One half was used as an archive, whereas the other half were split into quarters for microbiological and for geochemical, sedimentological and micropalaeontological analyses.

### Molecular sample preparation and geochemical analysis

The core was sectioned at different intervals. The core of the Mamontov Klyk site (C2) was separated into 118 horizons that were used for pore water analyses. The Buor Khaya site (BK2) was divided into 80 horizons. After thawing of subsamples, pore water was sampled using Rhizones© with an effective pore diameter of 0.1 μm. The concentration of sulfate and nitrate was determined via a KOH eluent and a latex particle separation column on a Dionex DX-320 ion chromatograph. Manganese and iron were detected by a Perkin-Elmer ‘Inductively Coupled Plasma Optical Emission Spectrometry’ (ICP-OES) Optima 3000 XL.

For methane measurements, 3 g of frozen material were retrieved with ice screws and immediately immersed in 20 ml serum vials containing a saturated NaCl solution (315 g l^-1^). Serum vials were sealed with butyl-rubber stopper and a crimp seal. Headspace gas was measured in triplicates with different setups. In brief, gas was analyzed with an Agilent 6890 (19) or 7890A (13) gas chromatograph equipped with a flame ionization detector and with a carbon plot capillary column or HP-Plot Q (Porapak-Q) column. The temperature of the oven, injector and detector were 40, 120, and 160°C, respectively. In both cases, helium was used as carrier gas. The δ^13^C-CH_4_ were determined with an isotope ratio mass spectrometer (Finnigan Delta Plus) equipped with a PreCon and a GC/C III interface (Thermo, Bremen, Germany). The precision of replicated measurements was better than 0.5‰ VPDB. Methane δ^13^C-signatures were linked to the VPDB scale using internal (-43.8‰ VPDB) and external (RM8561; −73.27‰ VPDB) standards measured at least every 10 analyses.

### DNA extraction, 16S rRNA Illumina MiSeq sequencing and analysis

Genomic DNA of 4.7-13 g sediment of different depth (C2: 19 samples and BK2: 10 samples, table S2) was extracted after the protocol of Zhou *et al.*, 1996 (*71*). DNA concentrations were quantified with Nanophotometer® P360 (Implen GmbH, München, DE) and Qubit® 2.0 Flurometer (Thermo Fisher Scientific, Darmstadt, Germany) according to the manufacture’s protocols.

The 16S rRNA gene for bacteria was amplified with the primer combination S-D-Bact-0341-b-S-17 and S-D-Bact-0785-a-A-21. The 16S rRNA gene for archaea was amplified in a nested approach with the primer combination S-D-Arch-0020-a-S-19 and S-D-Arch-0958-a-A-19 in the first PCR for 40 cycles and S-D-Arch-0349-a-S-17 and S-D-Arch-0786-a-A-20 in the second PCR for 35 cycles, respectively. The primers were labelled with different combinations of barcodes that are listed together with primer sequences in Tables S4 and S5. The PCR mix contained 1x PCR buffer (Tris•Cl, KCl, (NH_4_)_2_SO_4_, 15 mM MgCl_2_; pH 8.7) (QIAGEN, Hilden, Germany), 0.5 μM of each primer (Biomers, Ulm, Germany), 0.2 mM of each deoxynucleoside (Thermo Fisher Scientific, Darmstadt, Germany), and 0.025 U μl^-1^ hot start polymerase (QIAGEN, Hilden, Germany). The thermocycler conditions were 95°C for 5 minutes (denaturation), followed by 40 cycles of 95°C for 1 minute (denaturation), 56°C for 45 seconds (annealing) and 72°C for 1 minute and 30 seconds (elongation), concluded with a final elongation step at 72°C for 10 minutes. PCR products were purified with a Hi Yield® Gel/PCR DNA fragment extraction kit (Süd-Laborbedarf, Gauting, Germany) according to the manufacture’s protocol. PCR products of 3 individual runs per sample were combined. PCR products of different samples were pooled in equimolar concentrations and compressed to a final volume 10 μl with a concentration of 200 ng μl^-1^ in a vacuum centrifuge Concentrator Plus (Eppendorf, Hamburg, Germany).

The sequencing was performed on an Illumina MiSeq sequencer (GATC Biotech, Konstanz) according to the standard protocol. The library was prepared with the MiSeq Reagent Kit V3 for 2x 300 bp paired-end reads according to the manufacture’s protocols. For better performance due to different sequencing length, we used 15% PhiX control v3 library.

The quality and taxonomic classification of the Illumina sequences were analyzed with the customized QIIME pipeline. For details see Supplementary material and methods.

### Construction of ANME-specific clone libraries

DNA of permafrost thaw horizons and methane peaks were investigated to amplify ANME communities with specific primers. Therefore we used an ANME-specific probe as reverse primer and a universal archaeal primer as forward primer (table S3). PCR mixes (25 μl) contained 1 x PCR buffer, 0.2 mM dNTP’s, 2 mM MgCl_2_, 0.08 mg ml^-1^ bovine serum albumin, and 0.02 units HotStart Taq Plus Polymerase (QIAGEN, Hilden, Germany) and were performed under the following conditions: initial denaturation 94 °C for 10 min, 30 cycles of denaturation 94 °C for 30 sec, annealing 59 °C for 1 min, extension 72 °C for 3 min, and a final elongation at 72 °C for 10 min. PCR amplicons were purified with the HiYield® Gel/PCR DNA Extraction Kit (SLG, Gauting, Germany) according to manufacturer’s protocols. Purified PCR products were cloned with the TOPO® TA Cloning® Kit (Invitrogen, Waltham, Massachusetts, USA) according to manufacturer’s protocols. Positive clones were directly sequenced by Sanger sequencing (GATC Biotech, Konstanz).

### Phylogenetic reconstruction

For phylogenetic analyses of 16S rRNA gene of archaeal and bacterial sequences, the ARB software package was used (72). After manual refinement of representative sequences of OTUs defined by the Illumina sequencing and partial sequences of clone libraries against the alignment of the SILVA 16S rRNA gene SSU reference database release 115 (73), phylogenetic trees were calculated. Phylogenetic reconstruction were based on the maximum-likelihood algorithm (PHYLIP-ML, 100 bootstraps) implemented in ARB with reference sequences (> 1400 bp for bacteria and > 850 bp for archaea) and the implement archaeal filter. Our own partial sequences were added to the tree using the maximum parsimony algorithm without allowing changes in tree topology.

### 454 pyrosequencing of functional genes and analysis

The *mcrA* fragment was amplified using the primer set mlas and mcrA-re (Table S4). In a touchdown PCR denaturation, annealing and elongation time was set at 1 min. The PCR conditions were as follows: initial denaturation at 95°C for 3 min, 15 cycles with a stepwise temperature decrement from 65°C to 50°C, followed by 15 cycles with an annealing temperature of 55°C and a final elongation at 72°C for 10 min. Tagging of amplicons with unique multiple identifier (MID) tags (Table S6) for 454 sequencing was conducted in a second PCR using amplicons from the touchdown PCR as template and 15 cycles with a constant annealing temperature of 55°C. PCR reactions were performed in several separate reactions and pooled till reaching at least 150 ng final product. We used the same chemicals as for 16S rRNA gene amplifications. PCR products were purified with a Hi Yield® Gel/PCR DNA fragment extraction kit (Süd-Laborbedarf, Gauting, Germany) according to the manufacture’s protocol. Amplicons were quantified with the Qubit 2.0 system with ds DNA HS assay kit (Invitrogen, Waltham, Massachusetts, USA), mixed in equimolar amounts and sequenced from both directions by Eurofins Genomics using Roche/454 GS FLX++ technology.

454 mcrA sequences were analyzed with the mothur software by a customized standard operating procedure. For details see Supplementary material and methods.

### Quantification of general functional genes

Quantitative PCR was performed using the CFX Connect™ Real-Time PCR Detection System (Bio-Rad Laboratories, Inc., Hercules, USA). Each reaction contained iTaq™ Universal SYBR® Green Supermix (12.5 μl per reaction of 2x concentrate, Bio-Rad Laboratories, Inc., Hercules, USA), PCR primers (0.5 μl containing 20 μM each), sterile water (6.5 μl) and DNA template (5 μl) added to a final volume of 25 μl. Primers targeting the general functional genes *mcrA* and *dsrB* were used (table S3). The PCR reactions comprised an initial denaturation (5 min at 95°C), followed by 40 cycles of 5 s at 95°C, 30 s at the specific annealing temperature (see table S3), 10s at 72°C and a plate read step at 80°C for 3 s. Melt curve analysis from 65 to 95°C with a 0.5°C temperature increment per 5 s cycle was conducted at the end of each run to identify non-specific amplification of DNA. The qPCR assays were calibrated using known amounts of PCR amplified and cloned gene fragments from corresponding taxa (standards of pure cultures) in the range of 10^6^ – 10^1^ copies μl^-1^. Prior to qPCR analysis, DNA templates were diluted 5- to 100-fold and 3 replicates were analyzed for each sample. The PCR efficiency based on the standard curve was calculated using the BioRad CFX Manager software and varied between 88 and 100%, depending on the standard. All cycle data were collected using the single threshold cq determination mode.

### Quantification of ANME-2d

The DNA was diluted 1:100 to prevent inhibition of amplification by environmental compounds, e.g. humic acids, and to keep the cq value of the samples within detectable limits. The abundance of ANME-2d archaea was quantified by qPCR with specific *mcrA* primers (table S3). The PCR mix consisted of 10 μl KAPA HiFi SYBR green mix (KAPA Biosystems), primers (0.04 μl containing 100 mM each), MgCl_2_ solution (0.5 μl of 50 mM), bovine serum albumin (BSA) (0.3 μl), sterile water (9.02 μl) and 1 μl DNA template. qPCR amplifications were performed by 10 min 98°C initialization; 40 cycles of 5 second denaturation at 95°C, 30 second of annealing at 60°C, 1 min of elongation at 72°C, 2 seconds at 82°C for fluorescence detection; and finally a melting curve from 55°C to 95°C with a 0.5°C increment every 5 seconds. For quantification, a tenfold dilution series of mcrA156F/mcrA345R product cloned into a pGEM-T easy plasmid of a known copy number provided by Annika Vaksmaa of Radboud University, Nijmegen, was used as a standard (48). The PCR product specificity was checked by melt analysis and compared to the standard with a melting temperature of 82 °C.

### Microbial lipid biomarker extraction

Different fractions of lipids were extracted from sediment of the SMTZ. For a detailed method description see supplementary material.

### Catalyzed reporter deposition fluorescence in situ hybridization (CARD-FISH)

For CARD-FISH, sediments were fixed for 1 h at room temperature with 4 % formaldehyde (Fluka, Taufkirchen, Germany). For amplification, a fluorescine-labelled tyramide was used. The protocol for CARD-FISH followed the description published earlier (74). For details see supplementary material and methods.

### Statistical analysis of archaeal community

Statistical analysis was performed with the PAST 3.14 software (75). Canonical correlation analysis (CCA) was performed with 7 environmental parameters (chloride, pH, conductivity, methane, sulfate, manganese, iron, and nitrate) as explanatory variables and relative abundance of archaeal genera to families. Explanatory variables were standardized by log10 transformation prior to computation, with the exception of pH and conductivity. The significance of canonical axes was tested via permutation computed for N = 999. Spearman correlation analysis was performed with the implemented tools in PAST 3.14.

## Acknowledgement

We thank Antje Eulenburg, Ute Bastian, Katja Hockun, Cornelia Karger, Maria Bade and Anke Saborowski for excellent laboratory support. We further thank Aleksandr Maslov (SB RAS, Mel’nikov Permafrost Institute, Yakutsk, Russia) who provided indispensable drilling expertise. We thank Tiksi Hydrobase staff members Viktor Bayderin, Viktor Dobrobaba, Sergey Kamarin, Valery Kulikov, Dmitry Mashkov, Dmitry Melnichenko, Aleksandr Safin, and Aleksandr Shiyan for their field support and Birgit Schwinge (Universität Hamburg) for her help with the methane isotope measurements. We further thank Aurèle Vuillemin for helpful discussion on the manuscript.

## Author contributions

M.W., P.O. and S.L. planned the research. P.O. and M.G. conducted the field work. M.W. J.M. and R.R. conducted the molecular work. M.W. and F.H. analyzed the sequences. P.O., M. Wf. conducted the chemical measurements. C.K. analyzed the isotopic measurements of methane. M.W. and K.M. performed lipid analysis. M.W. carried out statistics. D.W. facilitated laboratory work and contributed to the discussion; M.W. and S.L. wrote the manuscript with input from all authors.

## Data deposition

Sequences of the submarine permafrost metagenome have been deposited at the NCBI Sequence Read Archive (SRA) with the Project number BioProject ID# PRJNA352907, under the accession numbers SRR5183846-SRR5183871 for archaeal 16S rRNA gene sequences, SRR5184420-SRR5184446 for bacterial 16S rRNA gene sequences, and SRR5019811-SRR5019818 for *mcrA* sequences. ANME-specific, partial 16S rRNA gene sequences of clone libraries are available in Genbank, EMBL and DDBJ under the accession numbers KY613956-KY613991.

